# Comparative validation of breast cancer risk prediction models and projections for future risk stratification

**DOI:** 10.1101/440347

**Authors:** Parichoy Pal Choudhury, Amber N. Wilcox, Mark N. Brook, Yan Zhang, Thomas Ahearn, Nick Orr, Penny Coulson, Minouk J. Schoemaker, Michael E. Jones, Mitchell H. Gail, Anthony J. Swerdlow, Nilanjan Chatterjee, Montserrat Garcia-Closas

## Abstract

**Background:** Well-validated risk models are critical for risk stratified breast cancer prevention. We used the Individualized Coherent Absolute Risk Estimation (iCARE) tool for comparative model validation of five-year risk of invasive breast cancer in a prospective cohort, and to make projections for population risk stratification.

**Methods:** Performance of two recently developed models, iCARE-BPC3 and iCARE-Lit, were compared with two established models (BCRAT, IBIS) based on classical risk factors in a UK-based cohort of 64,874 women (863 cases) aged 35-74 years. Risk projections in US White non-Hispanic women aged 50-70 years were made to assess potential improvements in risk stratification by adding mammographic breast density (MD) and polygenic risk score (PRS).

**Results:** The best calibrated models were iCARE-Lit (expected to observed number of cases (E/O)=0.98 (95% confidence interval [CI]=0.87 to 1.11)) for women younger than 50 years; and iCARE-BPC3 (E/O=1.00 (0.93 to 1.09)) for women 50 years or older. Risk projections using iCARE-BPC3 indicated classical risk factors can identify ~500,000 women at moderate to high risk (>3% five-year risk). Additional information on MD and a PRS based on 172 variants is expected to increase this to ~3.6 million, and among them, ~155,000 invasive breast cancer cases are expected within five years.

**Conclusions:** iCARE models based on classical risk factors perform similarly or better than BCRAT or IBIS. Addition of MD and PRS can lead to substantial improvements in risk stratification. Independent prospective validation of integrated models is needed prior to clinical evaluation risk stratified breast cancer screening and prevention.

## Introduction

Breast cancer risk prediction models are used in clinical and research settings to identify women at elevated risk of disease who could benefit from preventive therapies, enhanced screening, or be eligible to participate in prevention trials. Continuing updates of risk models incorporating additional risk factors (e.g., mammographic breast density (MD) and newly discovered genetic variants) will potentially improve our ability to identify such women.^1^

Prospective validation of models in independent studies is critical to determine their accuracy of prediction, robustness and potential for clinical application. BCRAT and IBIS are two established models that originally included hormonal and environmental risk factors and are currently used for clinical and research applications.^2^ BCRAT has been extensively validated, generally showing good calibration but low risk discrimination.^2-4^ IBIS performed better than BCRAT in terms of calibration and discrimination in an average-to high-risk population.^5^ Addition of MD^6-10^ or polygenetic risk scores (PRS)^11-19^ can lead to improved risk stratification. However, prospective evaluation of the accuracy of breast cancer absolute risk predictions from models incorporating PRS is currently lacking. This is different from comparing risk scores between women with and without breast cancer at a single time point,^12,16,17^ that evaluates risk discrimination, but not the accuracy of the model.

It is important that risk models remain dynamic and flexible in their ability to incorporate information on additional risk factors and context-specific incidence rates. However, a major challenge with developing and validating a comprehensive model is that information on all relevant risk factors is often not available in a single study. It is increasingly possible to develop models integrating information from multiple epidemiologic studies and other data sources using novel methods for data integration.^20-22^ We have recently developed the Individualized Coherent Absolute Risk Estimation (iCARE) software, implementing a flexible approach to build models for absolute risk combining information on relative risk estimates, incidence/mortality rates and risk factor distributions, with the latter two for a given target population and time period, from multiple data sources.^23-25^ It includes advanced features to account for missing risk factors using internal imputation and a validation component to facilitate comparative evaluation of multiple models across multiple cohorts using uniform methodology.^25^

We have previously used iCARE to develop a breast cancer risk model using relative risks from a multivariate regression based on eight prospective cohorts of women aged 50 years or older (iCARE-BPC3).^23^ Here, we develop an updated version of the synthetic model, described in Garcia-Closas et al^1^, based on relative risks from published literature (iCARE-Lit). A literature-based model, while requiring more assumptions, can include comprehensive sets of risk factors that may not all be measured in one study.

The current study aims to compare the performances of the iCARE models, BCRAT and IBIS based on classical risk factors, obtained by self-report through a short questionnaire, in the UK-based Generations Study (GS). In addition, the Prostate, Lung, Colorectal and Ovarian (PLCO) Cancer Screening Trial, which was used to develop iCARE-BPC3, served for further evaluation of the other models. Risk projections in a target population were estimated based on classical risk factors, and after addition of MD^26^ and PRS.

## Materials and Methods

### Study populations

Data were used from two prospective cohorts: GS from the UK and PLCO from the US.^27,28^ 113,211 women from GS, aged 16-102 years at enrollment (2003-2012), were considered. Participants from PLCO were between ages 50-75 years at enrollment (1993-2001) (N=78,214 women). Exclusion criteria included history of breast cancer, non-white or unknown ethnicity, no genetic consent or source of DNA, and age at entry below 35 or above 75 years. GS subjects were further excluded if they had a first-or second-degree relative in study. PLCO subjects with unconfirmable report of breast cancer were further excluded. The final analytic samples from the GS and PLCO included 64,874 (863 cases within five years) and 48,279 (1,008 cases within five years), respectively (**Supplementary Figure 1**). As PLCO was used for the development of iCARE-BPC3,^23^ it was only used for validation of other models. **Supplementary Table 1** shows the risk factor distributions in both validation cohorts.

**Table 1.** Ratios of expected to observed five-year absolute risk for the breast cancer risk prediction models validated using the GS* * AUC = area under the curve; GS = Generations Study; CI = confidence interval; E = expected absolute risk; O = observed absolute risk. The AUCs reported in Table 1 are defined based on absolute risk and incorporate the variation due to age. The AUCs (95%CI) based on the relative risk score, which do not incorporate variation of age, are as follows: (i) for women younger than 50 years, iCARE-Lit: 58.8 (55.3 to 62.3), BCRAT: 54.6 (50.9 to 58.4), IBIS: 57.0 (53.4 to 60.6); (ii) for women 50 years or older, iCARE-Lit: 60.3 (58.0 to 62.6), iCARE-BPC3: 57.7 (55.4 to 60.0), BCRAT: 52.2 (49.6 to 54.7), IBIS: 60.2 (57.9 to 62.5).

| Age group | Model | AUC (95% CI) | Overall |  |  | Top risk decile |  |  |
| --- | --- | --- | --- | --- | --- | --- | --- | --- |
|  |  |  | O% (95% CI) | E% | E/O ratio (95% CI) | O% (95% CI) | E% | E/O ratio (95% CI) |
| <50 years of age<br>(265 cases, 27,967 non-cases) | iCARE-Lit | 65.4 (62.1 to 68.7) | 0.94 (0.83 to 1.05) | 0.92 | 0.98 (0.87 to 1.11) | 2.22 (1.68 to 2.77) | 2.51 | 1.13 (0.89 to 1.44) |
|  | BCRAT | 64.0 (60.6 to 67.4) |  | 0.79 | 0.85 (0.75 to 0.95) | 2.24 (1.70 to 2.79) | 1.73 | 0.77 (0.60 to 0.98) |
|  | IBIS | 64.6 (61.3 to 67.9) |  | 1.07 | 1.14 (1.01 to 1.29) | 2.19 (1.65 to 2.73) | 2.58 | 1.18 (0.92 to 1.51) |
| ≥50 years of age<br>(598 cases, 36,044 non-cases) | iCARE-Lit | 62.2 (60.0 to 64.5) | 1.63 (1.50 to 1.76) | 1.84 | 1.13 (1.04 to 1.22) | 3.20 (2.62 to 3.77) | 3.91 | 1.22 (1.02 to 1.46) |
|  | iCARE-BPC3 | 60.2 (58.0 to 62.4) |  | 1.64 | 1.00 (0.93 to 1.09) | 2.63 (2.12 to 3.15) | 2.85 | 1.08 (0.89 to 1.32) |
|  | BCRAT | 58.2 (55.8 to 60.5) |  | 1.56 | 0.95 (0.88 to 1.03) | 2.73 (2.20 to 3.26) | 3.27 | 1.20 (0.99 to 1.46) |
|  | IBIS | 61.4 (59.2 to 63.6) |  | 1.85 | 1.13 (1.05 to 1.23) | 3.07 (2.51 to 3.63) | 3.98 | 1.30 (1.08 to 1.56) |

### Breast Cancer Risk Model Validation and Risk Projection

The supplements provide detailed descriptions of iCARE-based models (**Supplementary Table 2, Supplementary Table 3**) as well as BCRAT and IBIS **(Supplementary Table 4).** All models incorporate information on marginal disease incidence rates (**Supplementary Figures 2A and 2B**) and account for competing causes of mortality using mortality rates, both available from population-based registries^25,29,30^. The incidence rates were used to calibrate the average predicted risk to the national breast cancer risk.^29,30^ iCARE implements this step using an additional individual-level reference dataset of risk factors representing the underlying population.

For evaluating calibration, we categorized individuals based on deciles of both the five-year absolute risks that incorporates the variation of age and relative risk-score (i.e., sum of log relative risks multiplied by risk factors) that does not incorporate age information. The predicted and observed risks across risk categories were compared using expected-to-observed ratio and goodness-of-fit tests. Model discrimination was assessed using the area under the curve (AUC) statistics based on both five-year absolute risk and the relative risk-score (details in supplements).

Risk projections of invasive breast cancer were estimated among US non-Hispanic White women aged 50-70 years using the best calibrated model based on classical risk factors in that age group. We explored potential improvements in risk stratification with addition of PRS and MD. Apart from the 172-SNP PRS,^31,32^ hypothetical improved versions of PRS were considered that capture in part or wholly the missing common variant heritability.^31,33^ Theoretical AUC was computed based on relative risk-scores for different combinations: classical risk factors only, PRS only, MD only, and a combined model with all risk factors. We also estimated the number of women and future cases identified at the extremes of the risk distribution (details in supplement).^23,34^

## Results

### Breast Cancer Risk Model Validation

Among women younger than 50 years, all models (iCARE-Lit, BCRAT and IBIS) showed good calibration of relative risk (**Supplementary Figure 3A**). Absolute risk was best calibrated for iCARE-Lit (E/O (95% CI)=0.98 (0.87 to 1.11)) with AUC (95% CI)=65.4 (62.1 to 68.7) (**Table 1**). BCRAT tended to underestimate (E/O (95% CI)=0.85 (0.75 to 0.95)) and IBIS to overestimate (E/O (95% CI)=1.14 (1.01, 1.29)) absolute risk. For both models, the miscalibration was most evident in the highest risk deciles (**Figure 1, Table 1**).

**Figure 1.**
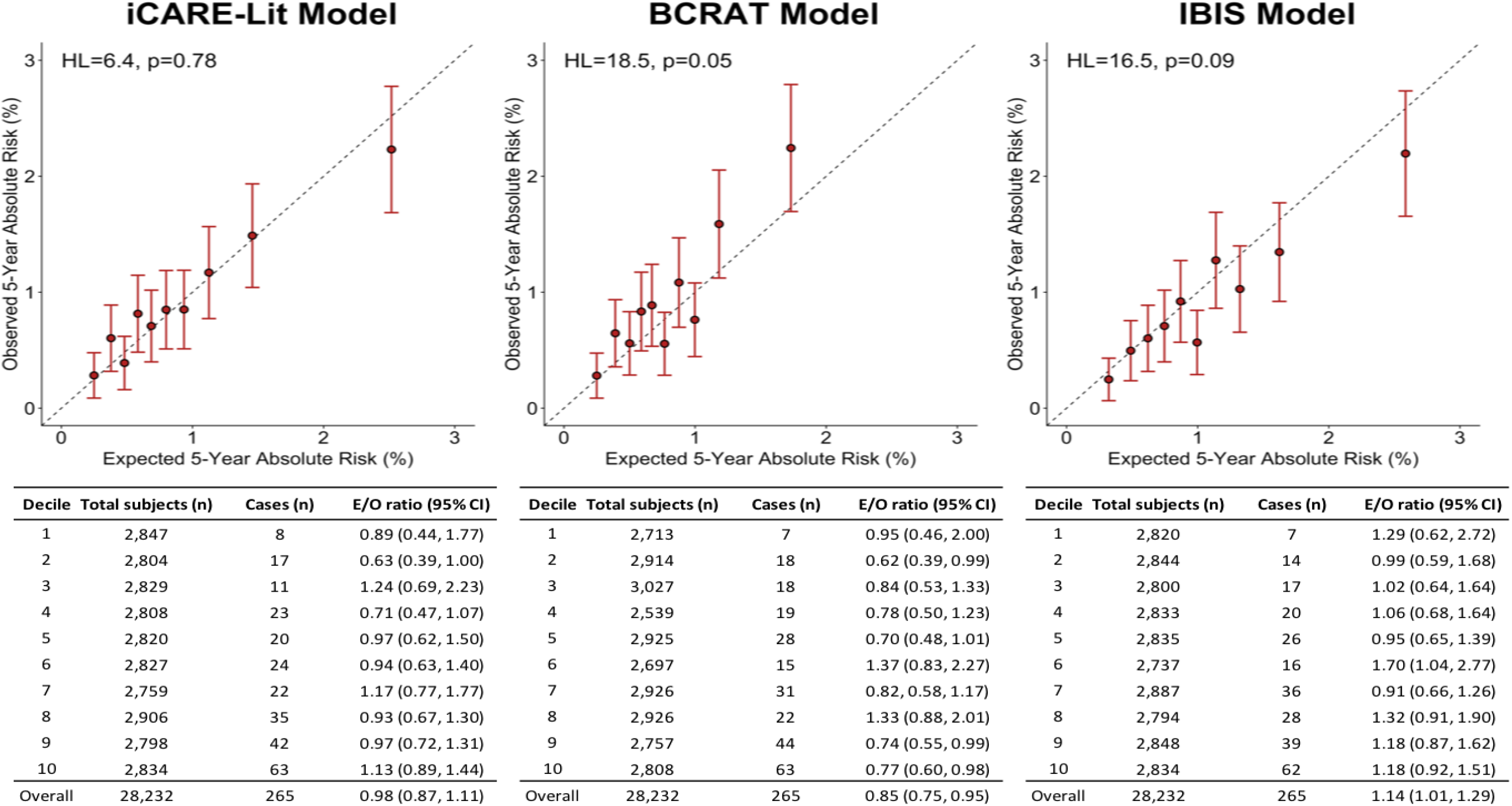
Absolute risk calibration of breast cancer risk prediction models in the GS cohort among women less than 50 years of age. The risk categories are based on absolute risk. GS = Generations Study; HL = Hosmer-Lemeshow test statistic.

Among women aged 50 years or older, iCARE-BPC3 showed good calibration of absolute and relative risk with E/O=1.00 (0.93 to 1.09) and AUC=60.2 (58.0 to 62.4). iCARE-Lit showed good calibration of relative risk but overestimation of absolute risk (E/O (95% CI)=1.13 (1.04 to 1.22)) (**Table 1, Figure 2, Supplementary Figure 4A**). BCRAT and IBIS showed miscalibration of both absolute and relative risk. While BCRAT was fairly well calibrated overall (E/O (95% CI)=0.95 (0.88 to 1.03)) with some underestimation in low-risk deciles and overestimation in high-risk deciles, IBIS (E/O (95% CI)=1.13 (1.05 to 1.23)) showed similar extent of overall miscalibration as iCARE-Lit. (**Table 1, Figure 2, Supplementary Figure 4A**).

**Figure 2.**
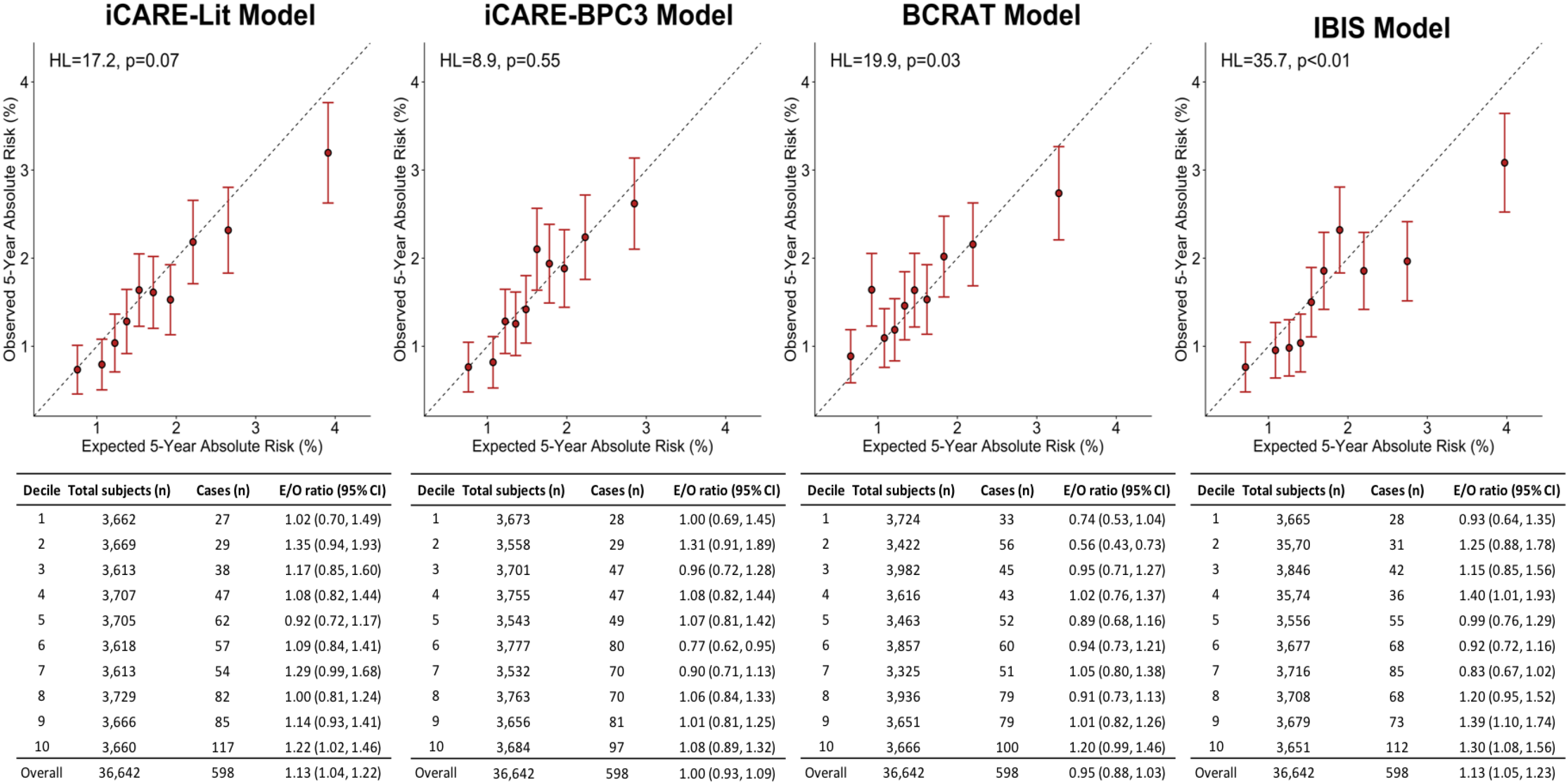
Absolute risk calibration of breast cancer risk prediction models in the GS cohort among women 50 years of age or greater. The risk categories are based on absolute risk. GS = Generations Study; HL = Hosmer-Lemeshow test statistic.

In PLCO, iCARE-Lit produced overestimation of five-year absolute risk to a similar extent as in the GS for women aged 50 years or older. Both BCRAT and IBIS underestimated absolute risk (**Supplementary Table 5, Supplementary Figure 5A**). In both cohorts, discriminatory accuracy was lower when risk categories were defined by the deciles of the relative risk-score, as opposed to absolute risk. (**Supplementary Figures 3B, 4B, 5B**).

### Breast Cancer Risk Projections

Five-year absolute risk projections based on the iCARE-BPC3 model were estimated among White non-Hispanic US women aged 50-70 years (~30 million women according to 2016 US Census). **Figure 3** shows the distribution of projected absolute risks and theoretical AUCs, based on relative risk-scores, for different risk factor combinations. Compared to classical risk factors, MD alone achieves an improved AUC. Further improvement is seen when including PRS alone. An integrated model with classical risk factors, MD and PRS gives the most improved AUC of 68.8 (**Figure 3**).

**Figure 3.**
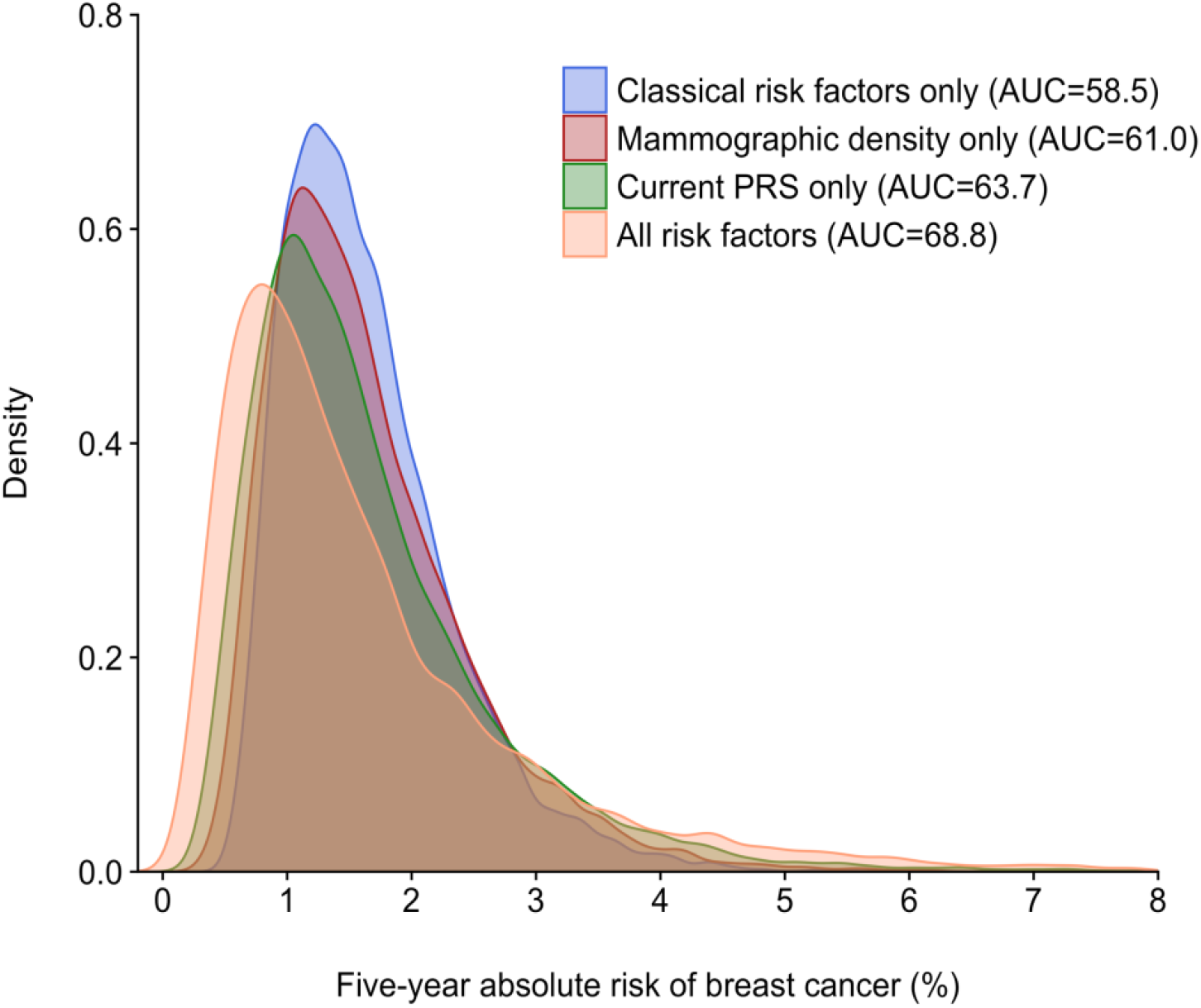
Five-year absolute risk projection for the general US population of White women, ages 50-70 years. The classical risk factors correspond to the iCARE-BPC3 model. AUC = area under the curve; PRS = polygenic risk score. The AUCs reported are based on the relative risk-score in that population and do not incorporate variation due to age.

Based on classical risk factors alone, approximately 2,400 women, representing <0.1% of the above population, could be identified at low risk (<0.5% five-year risk) of invasive breast cancer (**Figure 4B, Supplementary Table 6**). Only 11 of these women (<0.1% of all cases) are expected to develop the disease within five years. Additional information on MD and 172-SNP PRS is expected to increase the approximate number of women at low risk to 2.4 million, and around 8,800 of them (<2% of all cases) would be expected to develop the disease within five years. In the moderate-to high-risk group (>3% five-year risk), approximately 500,000 women, representing 1.7% of this population, would be identified based on classical risk factors, including approximately 17,000 (3.4% of all cases), expected to develop the disease within five years (**Figure 4C, Supplementary Table 6**). Adding MD and 172-SNP PRS increases the number of women identified to 3.6 million and among them, approximately 155,000 (~31% of all cases) would be expected to develop disease within five years.

**Figure 4.**
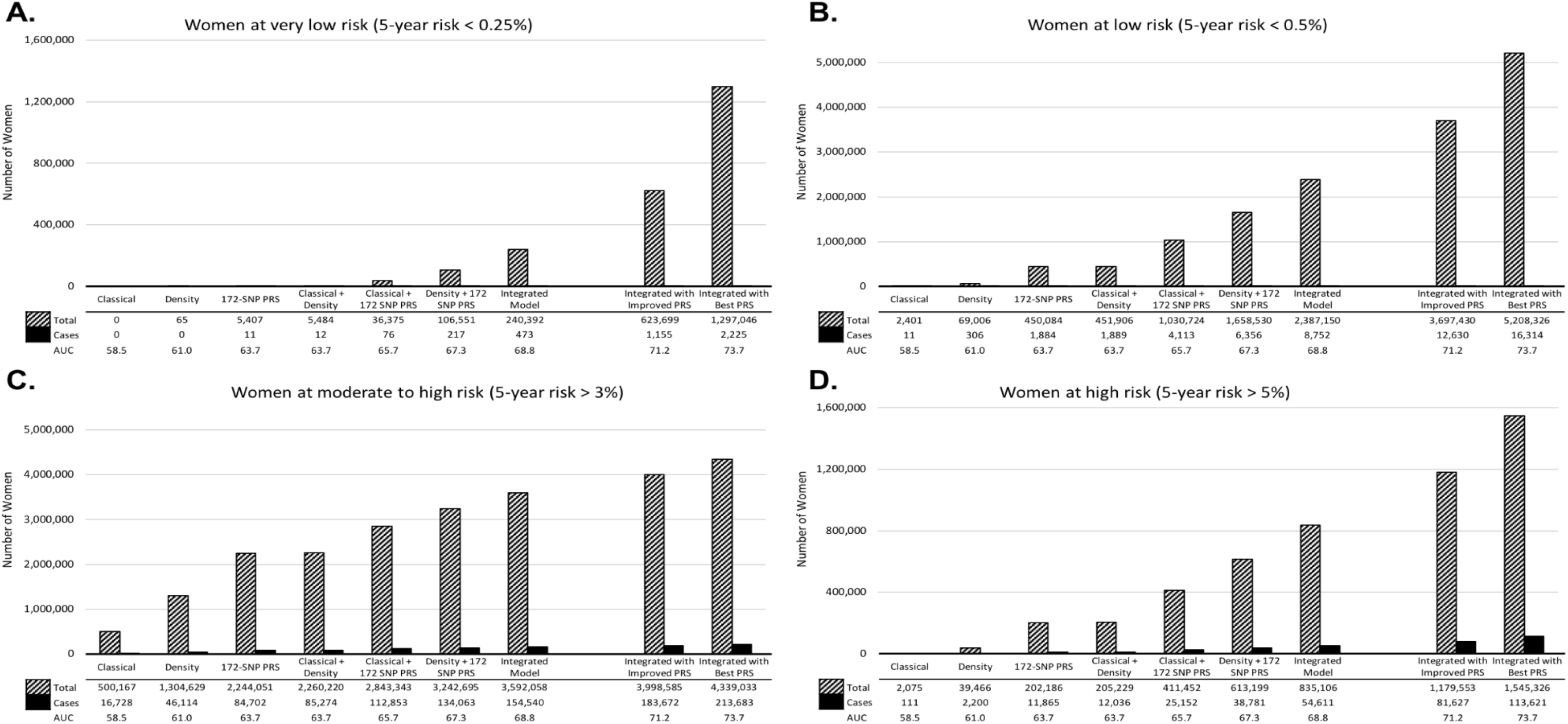
White non-Hispanic women aged 50-70 years in the US population expected to be identified at elevated risk of breast cancer according to different risk thresholds, and the incident cases of invasive breast cancer who are expected to occur in these groups within a five-year interval. The expected number of subjects is calculated using mid-2016 population estimates (n=30,030,821) from the US Census Bureau and the number of cases is calculated using an average predicted five-year risk of 1.61% and the 2015 invasive breast cancer incidence rates from SEER. AUC = area under the curve; PRS = polygenic risk score; SNP = single nucleotide polymorphism. The AUCs reported are based on the relative risk-score in that population and do not incorporate variation due to age.

We projected that doubling the size of current breast cancer GWAS (to around 300,000 cases and 300,000 controls) would yield additional discoveries and an improved PRS with AUC=67.3 (**Supplementary Table 6**). An integrated model with improved PRS could identify approximately 3.7 million women at low risk, and approximately 12,600 of these (2.6% of all cases) would be expected to develop invasive breast cancer within five years (**Figure 4B, Supplementary Table 6**). In the moderate-to high-risk group, we could identify close to 4 million women, and approximately 184,000 (37.3% of all cases), expected to develop the disease within five years. This is close to the risk stratification attained by the best theoretical PRS explaining 100% of the variability of polygenic risk from common variants (**Figure 4C, Supplementary Table 6**).

The relative increase in the numbers of women and cases identified due to incorporation of the improved PRS increases as the risk threshold becomes more extreme at either end of the distribution. For example, when comparing the integrated model with 172-SNP PRS to the integrated model with improved PRS, 11% more women and 19% more cases are identified above the 3% risk threshold, while 41% more women and 49% more cases are identified above the 5% risk threshold. On a similar note, the integrated model with improved PRS identified 54% more women below 0.5% risk threshold as opposed to 159% more women below 0.25% risk threshold. **(Figures 4A, 4B, 4C, 4D Supplementary Table 6)**.

## Discussion

In this comparative analysis using data from a large population-based cohort in the UK, we showed that iCARE-based absolute risk models for invasive breast cancer with classical risk factors are well calibrated, and that addition of MD and PRS can substantially improves risk stratification.

Among women younger than 50 years, all models were well calibrated in terms of relative risk; however, iCARE-Lit was most accurate for five-year absolute risk prediction. The finding that absolute risks may be miscalibrated while relative risks are not, illustrates the challenges of validating models for absolute risk, which requires incidence rates and the reference distribution of risk factors, ideally from the same time period as the validation study.

Among women aged 50 years or older, iCARE-BPC3 was well calibrated in terms of relative and absolute risk, while the other models overestimated absolute risk for women in the highest risk category. While relative risk is well calibrated for the iCARE-based models, the better absolute risk calibration for iCARE-BPC3 compared to iCARE-Lit could be due to the differences in specification of risk factors (i.e., finer versus coarser categories of continuous risk factors).

The IBIS and iCARE models include more information on classical risk factors than BCRAT,^29,30^ which has lower discriminatory accuracy. A distinct feature of IBIS is the modeling of detailed family history of cancer, considered important for identifying the women expected to be at very high risk due to extensive family history. We could not evaluate this potential advantage of IBIS due to miscalibration observed among women at higher risk.

The BCRAT has been extensively evaluated in previous studies and is currently recommended for predicting breast cancer risk for US women undergoing mammographic screening.^35-37^ BCRAT was well calibrated in some^38-40^ but not all validation studies, as several reported underestimation^41-43^ or overestimation^44-46^ of risk. Some studies reported improved model calibration on using incidence and mortality rates from the same country and time period.^42,43,46^ Sensitivity analyses using rates closer to our validation population also indicated slight improvement in calibration. Miscalibration may be due to model misspecification or differences in the risk factor distribution between the cohort and the underlying population.

The IBIS model was well calibrated in high-risk populations in the US and UK^5,47,48^. The IBIS 10-year risk was accurate as opposed to reported underestimates by BCRAT in a US-based study of average-to-high risk women.^5^ Both IBIS and BCRAT showed good absolute risk calibration in an Australian population of average-risk women.^15^ A recent prospective evaluation in a US-based integrated healthcare system showed good calibration of IBIS 10-year absolute risks overall, but ~20% overestimation in the highest risk decile (7.0% expected versus 5.9% observed).^8^ The current literature and our findings highlight the importance of prospective validation of risk models using multiple studies to evaluate robustness of model performance across populations.

Our study has several strengths. The GS is a recent population-based cohort that includes women with a wide age range. Moreover, the validation results of the iCARE-Lit model in the US-based PLCO study further supported our overall conclusions. We evaluated model calibration overall and stratified by levels of risk. The latter is important as accurate classification of subjects at the extremes of risk is most relevant for risk-based prevention and screening. Second, we assessed model calibration by deciles of expected absolute risk, as well as the relative risk-score. The former is commonly used in validation studies^5,38-41,46,47^ as absolute risk is the relevant measure for clinical or public health applications. However, strong dependence of absolute risk on age makes the differences in model performance due to risk factors other than age less evident than comparisons using the relative risk-score, which does not depend on age. We evaluated model calibration and discrimination with and without accounting for age. Most^5,39,41-44,47,48^ but not all^38,40,46^ previous validation studies of BCRAT and IBIS assessed model discrimination accounting for age. Such model discrimination statistics (e.g., AUC) evaluated in a validation cohort may differ from those in the target population owing to differences in risk-factor distributions. Moreover, our evaluation of discrimination at the extremes of the risk distribution showed that small changes in overall measures (e.g., AUC) derived from additional risk factors can result in substantial changes in the number of women that cross high-or low-risk thresholds with potential impact screening and prevention strategies.

Limitations of our analyses include that not all risk factors were available in the validation cohort. Although we used two studies to assess model performance, additional validation in independent cohorts, preferably embedded in representative health care systems where models will be applied, is desirable for more robust model evaluation. Moreover, we only evaluated short-term risk predictions, assuming risk factors remaining constant over the prediction period. Validation of long-term risk will require further follow up or additional studies, preferably accounting for time-varying risk factors and time-dependent associations.

Evaluation of potential impact of risk stratified prevention strategies requires estimation of risk stratification in a target population, as opposed to a particular validation cohort. The US Preventive Services Task Force recommends that women at moderate-to high-risk assess their risk-benefit trade-off of using chemoprevention with a healthcare professional.^49^ Although use of selective estrogen receptor modulators could result in up to 40% reduction of breast cancer risk in women at elevated risk,^50^ exploring alternative formulations with lower side effects may increase acceptability.^51^ Moreover, identification of women with low breast cancer risk who could benefit from less frequent screening will potentially improve the cost-effectiveness of screening programs by reducing overdiagnosis while preserving its potential benefits.^19^

Our projections based on an integrated model assumed that risk factors, MD (adjusted for age and BMI) and PRS act multiplicatively on disease risk. While we accounted for known dependencies between the risk factors and previous studies support multiplicative effects of risk factors on disease risk,^23,52^ these integrated models require empirical validation in independent prospective studies. We derived PRS based on SNP odds ratios from genetic discovery studies.^31,32^ This may lead to overestimation of risk stratification; however, based on previous assessment^53^ this bias is likely to be small.

Updates to BCRAT and IBIS have added MD^6-10^ and PRS^11-17^ to their original models; however, only the addition of MD to IBIS has been prospectively evaluated.^8,17^ This study demonstrated the value of aggregating information from classical risk factors and MD to identify women at elevated risk of breast cancer, but risk for women in the highest risk decile was overestimated.^8^ Addition of a 18-SNP PRS to IBIS was shown to increase risk discrimination in a UK-based screening cohort, though accuracy of prediction was not prospectively evaluated.^17^ The Breast Cancer Surveillance Consortium risk model, which includes BI-RADS MD and a 76-SNP PRS, was also not validated prospectively.^54^

Further improvements in risk stratification could be achieved by incorporating additional risk factors and heterogeneity in risk factor associations by breast cancer subtypes. Ultimately, it is desirable to develop a comprehensive model, robustly validated in multiple populations, applicable in populations with a wide range of underlying risk, and capable of handling missing risk factors to provide reliable risk estimates based on subsets of risk factors depending on the clinical application (e.g., risk assessment before or after mammography). The iCARE methodology facilitates this goal by providing a flexible risk modeling and validation tool.

In conclusion, we have demonstrated that iCARE models based on classical risk factors perform similarly and, in some cases, better than established models for five-year risk predictions of invasive breast cancer. Substantial improvements in risk stratification can be achieved with addition of MD and PRS to classical risk factors. Further development and evaluation of models in other racial/ethnic groups, and empirical validation of integrated risk models in prospective cohort studies is needed prior to broad clinical or research applications.

## Funding

This work was supported by the Intramural Research Program of the National Institutes of Health, NCI, Division of Cancer Epidemiology and Genetics (Z01CP010119), and the European Union’s Horizon 2020 research and innovation programme under grant agreements No 633784 (B-CAST).

The Generations Study would like to thank Breast Cancer Now and the Institute of Cancer Research for support and funding of the Generations Study, and the Study participants, Study staff, and the doctors, nurses and other health care staff and data providers who have contributed to the Study. The ICR acknowledge NHS funding to the NIHR Biomedical Research Centre.

## Notes

We would like to thank Adam R. Brentnall and Jack Cuzick for providing the IBIS Risk Evaluator software and for their comments to a draft of the manuscript.

We would like to thank Celine Vachon for providing unpublished information on the association of mammographic breast density with invasive breast cancer and population distribution of mammographic density. This information was used in the risk projection calculations.

## SUPPLEMENTARY MATERIALS

### 1.0 Cohort Populations for Validation

#### 1.1 Definition of Follow-Up

The Generations Study (GS) subjects received their first and second follow-up questionnaires approximately 2.5 years (median=2.53, IQR=2.48, 2.59) and 6 years (median=6.13, IQR=5.91, 6.46) after study entry, respectively. In addition to risk factors, the questionnaires sought information on breast cancer diagnosis.^1^ Breast cancer reports were confirmed by cancer registry, hospital or pathology records. Follow-up was defined from the date of study entry to the date the latest of the follow-up questionnaires was due. A small fraction of the participants (2.9%) were lost to questionnaire follow-up and most of them (~97%) agreed to be flagged with the National Health Service Central Registers to determine breast cancer and vital status. These participants were censored at June 30, 2012 because (at the time of data preparation) national cancer incidence data were incomplete after that date; the remaining participants were censored at loss to follow-up.

In the Prostate, Lung, Colorectal and Ovarian (PLCO) Cancer Screening Trial cohort (PLCO) subjects were followed up annually by the recruitment centers to ascertain cancer diagnosis status until trial year 13 or until December 31^st^ 2009, whichever came first. Analyses included women from both the control and intervention arms of PLCO. Sensitivity analyses restricted to the control arm yield similar results, therefore, we only provide full cohort analyses.

#### 1.2 Age Range of the Validation Cohorts

Subjects less than 35 years of age at entry were excluded because this is the minimum age for the BCRAT model and subjects greater than 75 years of age at entry were excluded so that five-year risk predictions are not made beyond 80 years of age.

#### 1.3 Risk Factors in the Validation Cohorts

There are several risk factors in the IBIS model for which the two validation cohorts in this analysis (GS and PLCO) did not have data: benign breast disease (BBD) pathology, Ashkenazi inheritance, personal BRCA 1/2 status and status of relatives, bilateral breast cancer (status and age), number of nieces, and data on cousins. Additionally, the GS did not have information on personal history of ovarian cancer, as well as ovarian cancer status and breast or ovarian cancer age among aunts. PLCO was additionally missing information on (hormone replacement therapy) HRT use, as well as data on grandmothers and aunts.

Due to certain limitations of the PLCO questionnaires, some assumptions were made when performing the IBIS model analysis in this cohort. For subjects with more than one relative of a particular type (i.e., sister or daughter), the questionnaire provided a summary count of the total number of each type of cancer (breast and ovarian) among the relatives. In such situations, we assumed that there were multiple relatives, each with a cancer, rather than multiple cancers in a single relative. In addition, the baseline questionnaire, completed between 1993-2001, provided an aggregate count for number of sisters (not distinguishing between half and full). Therefore, number of full sisters was ascertained from the supplemental questionnaire, completed between 2006-2008.

For the BCRAT model, both the GS and PLCO are missing data on history of atypical hyperplasia. PLCO is missing data on number of breast biopsies, so we assume that subjects with a history of BBD have had one breast biopsy, otherwise we assume no breast biopsies.

PLCO is missing data on HRT type among current users for the iCARE-Lit model and HRT type among ever users for the iCARE-BPC3 model.

### 2.0 Breast Cancer Risk Models

Below, we describe the breast cancer risk models evaluated in this analysis: iCARE-Lit and iCARE-BPC3 models and two established models (BCRAT and IBIS). **Supplementary Table 1** shows the risk factors included in each model.

#### 2.1 iCARE-Based Risk Models

We developed the iCARE-Lit model by combining estimates of relative risk parameters obtained from a literature review. The iCARE-Lit model has two sets of risk estimates and risk factor distributions: one for subjects less than 50 years of age and another for subjects 50 years of age or greater (**Supplementary Table 2**). This age stratification accounts for the fact that the relative risks for some risk factors (e.g., BMI and family history) are modified by age or menopausal status. This also accounts for the age-dependent distribution of some risk factors (e.g., parity and oral contraceptive use).

There are several differences in the specification of the iCARE-Lit model for women less than 50 years of age compared to the iCARE-Lit model for women 50 years of age or greater. iCARE-Lit (<50) includes two oral contraceptive (OC) use variables (never versus ever, current versus former or never) in order to be more flexible for validation cohorts that only have data on never/ever use. When there is no information on whether the ever OC user is a current or former user, this information is imputed using the reference dataset. The iCARE-Lit (≥50) model includes only never/ever OC use, assuming that all ever users are former users. iCARE-Lit (≥50) includes age at menopause and HRT use, while the iCARE-Lit (<50) model does not include these risk factors. Additionally, the relative risks for BBD, BMI, and family history of breast cancer vary between the iCARE-Lit (<50) and iCARE-Lit (≥50) models.

We previously developed the iCARE-BPC3 model for predicting absolute risk of breast cancer for White women in the U.S. 50 years of age or greater.2 The model was derived based on relative-risk parameters estimated from cohort studies participating in the Breast and Prostate Cancer Cohort Consortium (BPC3) (17,171 cases and 19,862 controls).

To predict risk for the UK-based cohort (GS), in both models we used incidence rates for invasive breast cancer and competing mortality rates from the UK Office for National Statistics (2006).^3,4^ Similarly, to predict risk for the US-based cohort (PLCO), we used incidence rates for invasive breast cancer from the US National Cancer Institute-Surveillance, Epidemiology, and End Results Program (NCI-SEER) (2008-2012)^5^ and competing mortality rates from the Center for Disease Control (CDC) WONDER database (2008-2012).^6^ We used information from various population-based surveys for estimating distribution of risk factors in both models. **Supplementary Table 2** and **Supplementary Table 3** describe the sources of information for the distribution of risk factors that are used to represent the UK and US populations.

#### 2.2 BCRAT Risk Model

The Breast Cancer Risk Assessment Tool (BCRAT), aka “Gail model”, was first developed in 1989 using data from a case-control study (2,852 cases and 3,146 controls) nested in the Breast Cancer Detection and Demonstration Project (BCDDP), a cohort of women undergoing annual screening mammogram in the US.^7^ The original model was subsequently modified in 1999 to project risk of developing invasive breast cancer using age-specific incidence rates from NCI-SEER and attributable risk estimates based on Cancer and Steroid Hormone (CASH) Study (https://seer.cancer.gov/archive/studies/epidemiology/study14.html), and the available algorithm used in this report was last updated in 2011 (http://www.cancer.gov/bcrisktool/).^8^ Relative risk scores and absolute risk estimates were obtained from the BCRAT Macro Version 3.0 (May 2011). Predictions for both validation cohorts were calculated using the BCRAT default incidence rates (i.e., SEER 1983-87 incidence rates).

#### 2.3 IBIS Risk Model

The International Breast Cancer Intervention Study model (IBIS version 8, aka “Tyrer–Cuzick Model”, http://www.ems-trials.org/riskevaluator/) uses family history information to compute a woman’s likelihood of carrying genes predisposing to breast cancer (in particular, BRCA1, BRCA2 and an additional low penetrance gene).^9^ The likelihood of carrying these genes is then used in conjunction with information on classical risk factors to estimate the probability of developing breast cancer over a specified period of time. We calculated expected five-year risk for both validation cohorts using software provided by the developers, which uses 2008-2010 breast cancer incidence and competing mortality rates obtained from Cancer Research UK.

### 3.0 Derivation of the Reference Populations

In order to project the distribution of absolute risk for the US and UK populations, we generated risk factor distributions that are representative of each population. These risk factor distributions were each coded according to the iCARE-BPC3 and iCARE-Lit models. Risk factor distributions are specified separately for women less than 50 years of age and women 50 years of age or older. These age categories are used as surrogates for menopausal status: we assume that women less than 50 years of age are premenopausal and never users of HRT, and women 50 years of age or older are postmenopausal and not current users of OC.

#### 3.1 US Reference Population

The US reference population is adapted from the reference population developed by Maas et al.^2^ The majority of the risk factors were derived from the 2008, 2010, and 2012 National Health and Nutrition Examination Survey (NHANES). An imputation model was used to simulate the distribution of alcohol intake based on the distribution among the controls in the Women’s Health Initiative (WHI) study. The imputation model was conditional on all variables in the iCARE-BPC3 model with significant associations with alcohol intake. The general strategy for imputation was to transform the variable to be normally distributed and then model the transformed variable conditional on other variables. At first the model was considered based on the regression of the transformed outcome based on each predictor separately to explore if there was a statistically significant association. In the second step, a joint model was fit that included all the predictors that were significantly associated with the transformed alcohol intake variable when considered individually. In this joint model, some of the predictors did not remain statistically significant and they were subsequently dropped one at a time starting with the one with the largest p-value. At each step likelihood ratio tests were used to compare each model with the reduced model after dropping the variable. After this process, the final model was used to impute the transformed outcome and that was subsequently back-transformed to get the actual outcome. A second model was used to determine whether the subject was a non-drinker. For subjects who were predicted to be non-drinkers, their alcohol consumption was set to zero, over-writing the previous imputation process. In the final step, the alcohol intake for each referent subject was imputed as an average predicted value plus a sampled value from the model residuals. The distribution of the imputed variable was close to the empirical data distribution. The distributions of the remaining risk factors were derived from the 2010 National Health Interview Survey (NHIS), the Prostate, Lung, Colorectal and Ovarian (PLCO) Cancer Screening Trial, and published literature.

#### 3.2 UK Reference Population

The risk factor distribution representative of the general UK population was simulated based on population-based surveys, cohort studies, and the published literature. The majority of variables were derived from the Health Survey for England (HSE) of 2005-2006. To account for missing data on OC and HRT status, the joint distribution was simulated for women 50 years of age or greater based on the conditional probabilities among those with data on these variables. In doing so, we are assuming that the status of HRT and OC use are missing at random.

In order to account for the correlation between parity and age at first birth, the joint distribution was derived from the Cohort Fertility Tables, England and Wales. In order to capture temporal affects in parity and age at first birth, the frequencies of these variables were averaged among women born between 1930 and 1955 to represent those aged 50 years or greater in 2006, and among women born between 1960 and 1975 to represent those less than 50 years of age in 2006. The contribution of each age interval from the fertility tables to the overall averages was weighted based on the age distribution of participants in the HSE surveys.

Multiple data sources were used to account for the correlation between body mass index (BMI) and age at menarche. Since the distribution of BMI in the GS would not be representative of the general UK population, the joint distribution of these two variables was simulated from the distribution of BMI in HSE, the distribution of age at menarche in the GS, and covariance between BMI and age at menarche in the GS.

The distributions of the remaining risk factors were derived from published literature.

### 4.0 Validation of Breast Cancer Risk Models

For each subject in the validation cohorts, person-time was accrued from the time of recruitment until the time of last contact or linkage with the subject. The follow-up time for each person for five-year risk prediction was defined from time of entry to five years after entry, time of last contact or linkage to cancer or death registries, whichever came first. The absolute risk of a woman developing breast cancer over the follow-up period was calculated accounting for competing risk due to death from other causes. From each risk prediction model (except IBIS), we obtained the relative risk score (i.e., linear predictor associated with the risk factors except age) and the expected absolute risk over five years since study entry for each subject in the validation cohorts. As the IBIS software does not provide relative risk scores, they were approximated as the ratio of the predicted absolute risk over one year since study entry and the incidence rate of invasive breast cancer at age of study entry.

The study subjects were classified into categories of low to high risk based on (a) deciles of the expected absolute risk, (b) deciles of the relative risk score. Within each risk category, the observed absolute risk (i.e., observed proportion of cases) and the average of the expected absolute risks over five years were computed after adjusting for observed follow-up.^10^ The 95% Wald-based confidence intervals for the observed absolute risks were computed using an influence function-based variance estimator.^10^ For each category, we also computed the observed and expected relative risks, which were defined for each woman as the ratio of her five-year absolute risk (observed and expected) and the average five-year absolute risk (observed and expected) in the validation study. The 95% Wald-based confidence intervals for observed relative risk were computed using the asymptotic variance estimator.^10^

We evaluated the overall discriminatory ability of each model using the Area Under the Curve (AUC), which was estimated as the empirical proportion of case-control pairs in which the absolute risk (or relative risk score for age-adjusted AUC) for a case was higher than the absolute risk (or relative risk score for age-adjusted AUC) for a control. We also computed the 95% Wald-based confidence interval for AUC using the asymptotic variance formula derived by DeLong et al.^11^

### 5.0 Breast Cancer Risk Projection

As a first step, we simulated current ages based on the distribution of ages in the 2016 US Census estimates for women 50-70 years in the US. In the next step, we estimate the five-year absolute risk and relative risk scores using the iCARE-BPC3 model and information from simulated current ages, the risk factor distribution from the reference dataset, and 2015 invasive breast cancer incidence rates from SEER.^12^ We use the standard deviation of the relative risk score evaluated based on the risk factor distribution to calculate theoretical AUCs for the projected risks.^13^ The mid-2016 population estimates from the US Census Bureau are used to translate the proportion of the population and of future cases identified as exceeding a pre-specified absolute risk threshold into corresponding numbers of subjects. The theoretical AUC depicts the amount of overall risk stratification achieved by the various risk factor combinations. For the risk factors for which reference data were not available (e.g., PRS, mammographic density), the relative risk scores were simulated using published estimates of their population level summary statistics (C. Vachon, unpublished data, 2018).^14-16^ Mammographic density was defined in relation to percent density and the log-odds ratio was adjusted for age and BMI (C. Vachon, unpublished data, 2018). We explored three versions of PRS: (1) the “current” PRS based on 172 SNPs discovered to date by GWAS,^14,16^ (2) an “improved” PRS including additional SNPs expected to be discovered as current GWAS sample sizes double (i.e., approximately a total of 300,000 cases and 300,000 controls); (3) and the “maximum” PRS that could explain heritability for breast cancer associated with log-additive effects of all common variants.^17^ These calculations were done using estimates of number of underlying susceptibility SNPs and distribution of their effects, which we obtained through application of a novel method for analysis of effect-size distribution underlying GWAS.^17^ With the overall familial relative risk (FRR) of breast cancer being 2,^14^ the overall heritability in the frailty scale or log-odds scale is calculated using the formula: *h*^2^ = 2 log(*FRR*) = 2*LOG*2 = 1.39. The “current” PRS is based on the common susceptibility variants that explain 18% of the overall FRR. The heritability explained by the “current” PRS is 0.18× 1.39 = 0.25. The “improved” PRS includes the additional SNPs expected to be discovered as the current GWAS sample sizes double (i.e., approximately 300,000 cases and 300,000 controls). The associated heritability (0.4) is calculated using a projection formula^17^ that uses information from the estimated common variant effect size distribution using the summary level statistics and external LD information. The “maximum” PRS based on all the SNPs that can be reliably imputed using OncoArray explain 41% of the overall FRR. The heritability explained by the “maximum” PRS is 0.41 × 1.39 = 0.57.^14^ In the simulations, we assume that the PRS is independent of epidemiologic risk factors given family history. For projections based on risk factor combinations with PRS, family history and other risk factors, we account for the attenuation of the relative risk for family history due to the correlation between PRS and family history.

---

**Table S1.** Risk factor distributions in the validation cohorts* * Risk factor distributions are reported as n (%), unless otherwise specified. BBD = benign breast disease, GS = Generations Study, C-type = estrogen and progestogen combined, E-type = estrogen-only, HRT = hormone replacement therapy, OC = oral contraceptive, PLCO = Prostate, Lung, Colorectal and Ovarian Cancer Screening Trial.

| Breast cancer risk factor | GS |  | PLCO |
| --- | --- | --- | --- |
|  | Ages <50, N=28,232 | Ages ≥50, N=36,642 | Ages ≥50, N=48,279 |
| **Age at baseline, years** |  |  |  |
| Median (range) | 42 (35-49) | 58 (50-74) | 61 (50-75) |
| **Age at menarche, years** |  |  |  |
| ≤11 | 5,600 (22.4) | 7,775 (23.3) | 9,612 (19.9) |
| 12-13 | 12,836 (51.2) | 16,210 (48.7) | 26,326 (54.6) |
| 14-15 | 5,777 (23.1) | 8,255 (24.8) | 10,224 (21.2) |
| ≥16 | 834 (3.3) | 1,077 (3.2) | 2,024 (4.2) |
| Missing | 3,185 | 3,325 | 93 |
| **Parity** |  |  |  |
| Nulliparous | 5,677 (20.1) | 4,524 (12.4) | 4,149 (8.6) |
| 1 birth | 4,155 (14.7) | 3,710 (10.1) | 3,270 (6.8) |
| 2 births | 12,879 (45.7) | 1,824 (50.9) | 11,523 (23.9) |
| 3 births | 5,491 (19.5) | 9,734 (26.6) | 29,270 (60.7) |
| Missing | 30 | 50 | 67 |
| **Age at first birth, years (among parous women)** |  |  |  |
| <20 | 910 (4.0) | 2,077 (6.5) | 7,786 (17.7) |
| 20-24 | 4,490 (20.0) | 10,759 (33.6) | 23,291 (53.0) |
| 25-29 | 8,658 (38.5) | 13,461 (42.1) | 9,710 (22.1) |
| ≥30 | 8,435 (37.5) | 5,690 (17.8) | 3,127 (7.1) |
| Missing | 62 | 131 | 216 |
| **OC use** |  |  |  |
| Never | 1,624 (5.8) | 7,493 (20.5) | 20,868 (43.3) |
| Ever | 26,582 (94.2) | 29,100 (79.5) | 27,378 (56.8) |
| Missing | 26 | 49 | 33 |
| **Current OC use (among women age <50)** |  |  |  |
| Never | 1,624 (5.8) |  |  |
| Former | 22,897 (81.2) | N/A | N/A |
| Current | 3,685 (13.1) |  |  |
| Missing | 26 |  |  |
| **HRT use (among women age ≥50)** |  |  |  |
| Never |  | 17,878 (49.0) | 14,303 (29.8) |
| Former |  | 6,191 (34.0) | 7,459 (15.5) |
| Current | N/A | 12,414 (17.0) | 26,271 (54.7) |
| Missing |  | 159 | 246 |
| **Type of HRT use (among current users age ≥50)** |  |  |  |
| Current E-type |  | 2,976 (52.7) |  |
| Current C-type | N/A | 2,676 (47.4) | N/A |
| Missing |  | 539 |  |
| **Age at menopause, years (among women age ≥50)** |  |  |  |
| <40 |  | 419 (1.9) | 6,718 (14.0) |
| 40-44 |  | 1,482 (6.8) | 6,728 (14.0) |
| 45-49 | N/A | 5,171 (23.8) | 11,156 (23.3) |
| 50-54 |  | 11,563 (53.2) | 17,688 (36.9) |
| ≥55 |  | 3,094 (14.2) | 5,624 (11.7) |
| Missing |  | 14,913 | 365 |
| **Height, m** |  |  |  |
| Median (range) | 1.7 (1.2-2.0) | 1.6 (1.3-2.1) | 1.6 (1.2-2.0) |
| **Body mass index, kg/m²** |  |  |  |
| <25 | 15,653 (56.7) | 17,912 (50.0) | 19,390 (40.6) |
| ≥25 - <30 | 7,702 (27.9) | 12,525 (34.9) | 16,775 (35.1) |
| ≥30 | 4,241 (15.4) | 5,423 (15.1) | 11,592 (24.3) |
| Missing | 636 | 782 | 522 |
| **Alcohol, g/day** |  |  |  |
| None | 4,227 (15.0) | 7,359 (20.5) | 17 (0.1) |
| <5 | 3,718 (13.2) | 5,000 (13.9) | 17,767 (74.4) |
| 5-14 | 8,374 (29.7) | 10,173 (28.4) | 2,900 (12.2) |
| 15-24 | 5,775 (20.5) | 7,032 (19.6) | 1,697 (7.1) |
| 25-34 | 2,948 (10.5) | 3,282 (9.1) | 405 (1.7) |
| 35-44 | 1,526 (5.4) | 1,533 (4.3) | 628 (2.6) |
| ≥45 | 1,621 (5.8) | 1,496 (4.2) | 460 (1.9) |
| Missing | 43 | 767 | 24,405 |
| **History of BBD** |  |  |  |
| No | 23,040 (81.6) | 26,669 (72.8) | 33,365 (70.2) |
| Yes | 5,192 (18.4) | 9,973 (27.2) | 14,136 (29.8) |
| Missing | 0 | 0 | 778 |
| **Breast cancer family history in first degree relatives** |  |  |  |
| No | 23,708 (84.0) | 30,528 (83.3) | 40,622 (85.6) |
| Yes | 4,524 (16.0) | 6,114 (16.7) | 6,837 (14.4) |
| Missing | 0 | 0 | 820 |

**Table S2.**
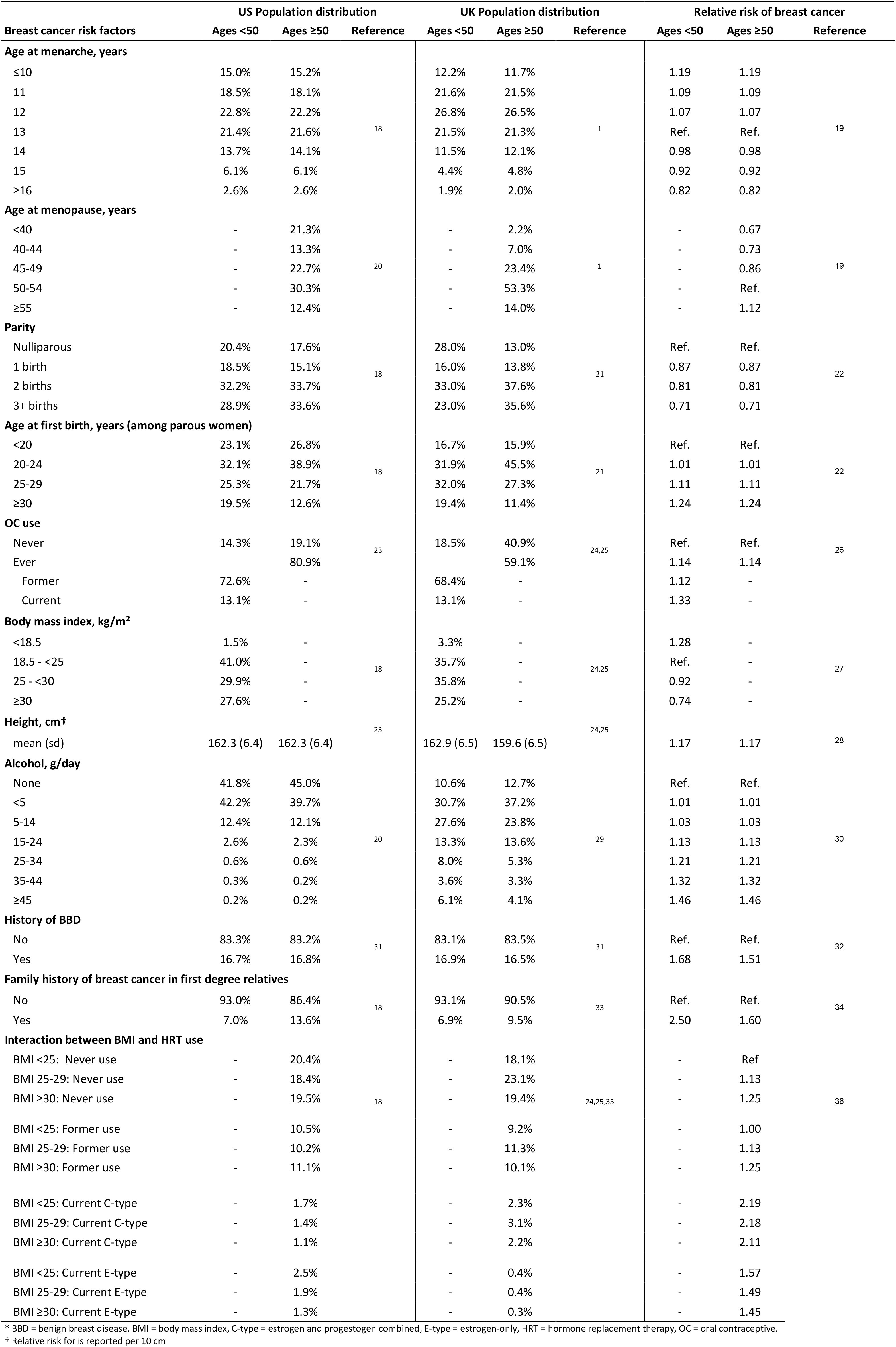
Parameters used for the development of the **iCARE-Lit** breast cancer risk prediction model* * BBD = benign breast disease, BMI = body mass index, C-type = estrogen and progestogen combined, E-type = estrogen-only, HRT = hormone replacement therapy, OC = oral contraceptive. † Relative risk for is reported per 10 cm

**Table S3.**
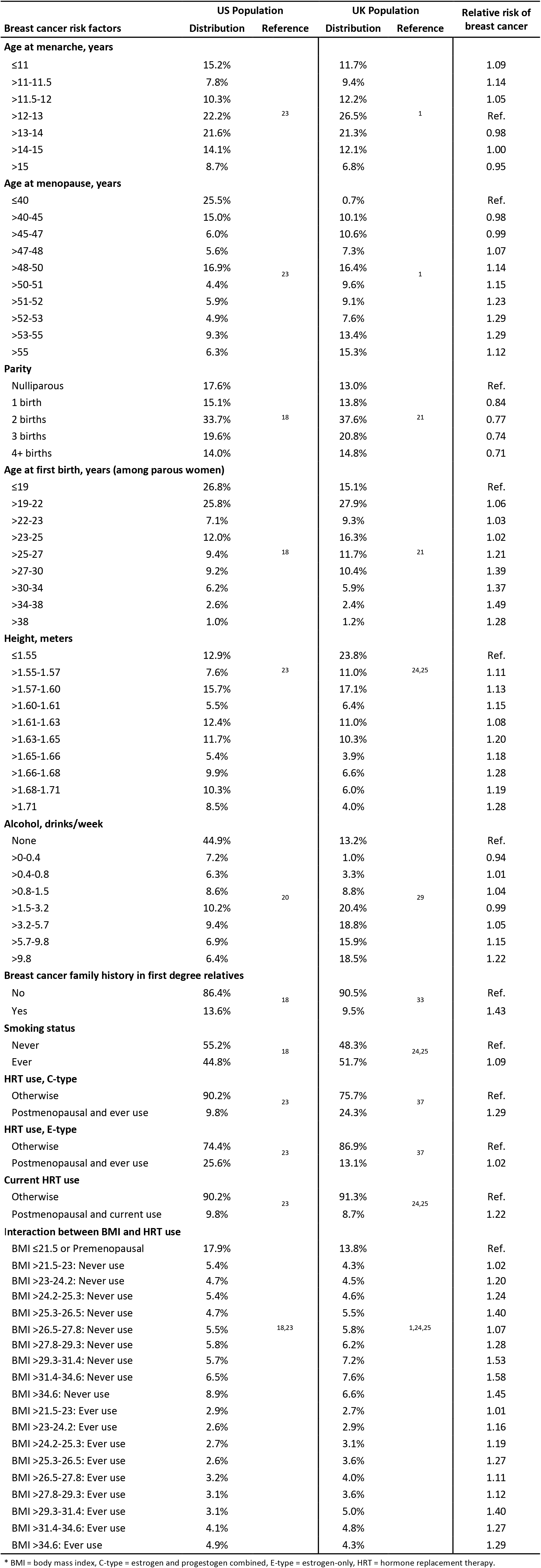
Parameters used for the development of the **iCARE-BPC3** breast cancer risk prediction model* * BMI = body mass index, C-type = estrogen and progestogen combined, E-type = estrogen-only, HRT = hormone replacement therapy.

**Table S4.** Risk factors included in the breast cancer risk prediction models* * BBD = benign breast disease. † iCARE-Lit and BCRAT use age 50 as a surrogate for menopausal status ‡ Assessed as yes/no for iCARE-Lit and as number of biopsies for BCRAT

| Risk factors included in the model | Breast cancer risk models |  |  |  |
| --- | --- | --- | --- | --- |
|  | iCARE-Lit | iCARE-BPC3 | BCRAT | IBIS |
| Age at menarche | ✓ | ✓ | ✓ | ✓ |
| Parity | ✓ | ✓ |  | ✓ |
| Age at first birth | ✓ | ✓ | ✓ | ✓ |
| Oral contraceptive use | ✓ |  |  |  |
| Body mass index (BMI) | ✓ | ✓ |  | ✓ |
| Alcohol use | ✓ | ✓ |  |  |
| Smoking status |  | ✓ |  |  |
| Menopausal status† |  | ✓ |  | ✓ |
| Age at menopause | ✓ | ✓ |  | ✓ |
| Menopausal replacement therapy use | ✓ | ✓ |  | ✓ |
| BBD‡ | ✓ |  | ✓ |  |
| Type of BBD |  |  |  |  |
| BBD - hyperplasia |  |  |  | ✓ |
| BBD - atypical hyperplasia |  |  | ✓ | ✓ |
| Lobular Carcinoma *in situ* (LCIS) |  |  |  | ✓ |
| Family history of breast and/or ovarian cancer |  |  |  |  |
| First-degree relatives with breast cancer (yes/no) | ✓ | ✓ | ✓ | ✓ |
| Number of first-degree relatives with breast cancer |  |  | ✓ | ✓ |
| Second-degree relatives with breast cancer |  |  |  | ✓ |
| Third-degree relatives with breast cancer |  |  |  | ✓ |
| Age of onset of breast cancer in a relative |  |  |  | ✓ |
| Bilateral breast cancer in a relative |  |  |  | ✓ |
| Ovarian cancer in a relative |  |  |  | ✓ |
| Male breast cancer |  |  |  | ✓ |

\* BBD = benign breast disease.
† iCARE-Lit and BCRAT use age 50 as a surrogate for menopausal status
‡ Assessed as yes/no for iCARE-Lit and as number of biopsies for BCRAT

**Table S5.** Ratios of expected to observed 5-year absolute risk for the breast cancer risk prediction models in PLCO (1,008 cases, 47,271 non-cases)* * AUC = area under the curve, CI = confidence interval, E = expected absolute risk, O = observed absolute risk; PLCO = Prostate, Lung, Colorectal and Ovarian Cancer Screening Trial. The AUCs reported in Supplementary Table 5 are defined based on absolute risk and incorporate the variation due to age. The AUCs (95%CI) based on the relative risk score, which do not include variation of age, are as follows: iCARE-Lit: 58.7 (57.0 to 60.5), BCRAT: 54.9 (53.0 to 56.8), IBIS: 58.1 (56.3 to 59.9).

| Model | AUC (95% CI) | Overall |  |  | Top risk decile |  |  |
| --- | --- | --- | --- | --- | --- | --- | --- |
|  |  | O% (95% CI) | E (%) | E/O ratio (95% CI) | O% (95% CI) | E (%) | E/O ratio (95% CI) |
| **iCARE-Lit** | 59.8 (58.1 to 61.6) |  | 2.46 | 1.18 (1.11 to 1.25) | 3.52 (3.00, 4.04) | 4.72 | 1.34 (1.16 to 1.55) |
| **BCRAT** | 57.2 (55.4 to 59.0) | 2.09 (1.96, 2.22) | 1.76 | 0.84 (0.79 to 0.90) | 3.14 (2.65, 3.63) | 3.81 | 1.21 (1.04 to 1.42) |
| **IBIS** | 58.8 (57.1 to 60.6) |  | 1.47 | 0.70 (0.66 to 0.75) | 3.36 (2.86, 3.87) | 3.02 | 0.90 (0.77 to 1.04) |

**Table S6.** Proportion of at-risk subjects and incident cases expected to be identified at different risk levels for models with different combinations of risk factors, among non-Hispanic White women ages 50-70 in the US* * The classical risk factors correspond to the iCARE-BPC3 model. The expected number of subjects is calculated using mid-2016 population estimates (n=30,030,821) from the US Census Bureau and the number of cases is calculated using an average predicted five-year risk of 1.64% and the 2015 invasive breast cancer incidence rates from SEER. AUC = area under the curve, AR = absolute risk, PRS = polygenic risk score, SNP = single nucleotide polymorphism. † PRS including additional SNPs expected to be discovered as current GWAS sample sizes double (i.e., approximately a total of 300,000 cases and 300,000 controls) ‡ PRS that could explain heritability for breast cancer associated with log-additive effects of all common variants § Odds ratio for mammographic breast density is adjusted for PRS The AUC is reported based on the relative risk-score in that population and do not incorporate variation of age.

| Incorporated breast cancer risk factors | AUC | Very low risk (5-year AR <0.25%) |  | Low risk (5-year AR <0.5%) |  | Moderate to high risk (5-year AR >3%) |  | High risk (5-year AR >5%) |  |
| --- | --- | --- | --- | --- | --- | --- | --- | --- | --- |
|  |  | Total subjects (n) | Cases (n) | Total subjects (n) | Cases (n) | Total subjects (n) | Cases (n) | Total subjects (n) | Cases (n) |
| Classical risk factors | 58.5 | <0.1% (0) | <0.1% (0) | <0.1% (2,401) | <0.1% (11) | 1.7% (500,167) | 3.4% (16,728) | <0.1% (2,075) | <0.1% (111) |
| Density | 61.0 | <0.1% (65) | <0.1% (0) | 0.2% (69,006) | 0.1% (306) | 4.3% (1,304,629) | 9.4% (46,114) | 0.1% (39,466) | 0.4% (2,200) |
| Current PRS (172-SNPs) | 63.7 | <0.1% (5,407) | <0.1% (11) | 1.5% (450,084) | 0.4% (1,884) | 7.5% (2,244,051) | 17.2% (84,702) | 0.7% (202,186) | 2.4% (11,865) |
| Classical risk factors + Density | 63.7 | <0.1% (5,484) | <0.1% (12) | 1.5% (451,906) | 0.4% (1,889) | 7.5% (2,260,220) | 17.3% (85,274) | 0.7% (205,229) | 2.4% (12,036) |
| Classical risk factors + Current PRS | 65.7 | 0.1% (36,375) | <0.1% (76) | 3.4% (1,030,724) | 0.8% (4,113) | 9.5% (2,843,343) | 22.9% (112,853) | 1.4% (411,452) | 5.1% (25,152) |
| Current PRS + Density | 67.3 | 0.4% (106,551) | <0.1% (217) | 5.5% (1,658,530) | 1.3% (6,356) | 10.8% (3,242,695) | 27.3% (134,063) | 2.0% (613,199) | 7.9% (38,781) |
| **All risk factors** | **68.8** | **0.8% (240,392)** | **0.1% (473)** | **7.9% (2,387,150)** | **1.8% (8,752)** | **12.0% (3,592,058)** | **31.4% (154,540)** | **2.8% (835,106)** | **11.1% (54,611)** |
| Improved PRS+ | 67.3 | 0.4% (108,387) | <0.1% (220) | 5.6% (1,667,306) | 1.3% (6,369) | 10.9% (3,272,719) | 27.5% (135,235) | 2.1% (624,603) | 8.0% (39,468) |
| Classical risk factors + Improved PRS+ | 68.8 | 0.8% (238,857) | 0.1% (469) | 7.9% (2,375,628) | 1.8% (8,693) | 12.0% (3,611,713) | 31.5% (155,143) | 2.8% (841,777) | 11.2% (54,951) |
| Improved PRS + Density†,§ | 69.9 | 1.3% (392,949) | 0.2% (752) | 10.0% (2,988,278) | 2.2% (10,609) | 12.6% (3,794,886) | 34.2% (168,416) | 3.3% (996,426) | 13.6% (66,930) |
| **All risk factors†,§** | **71.2** | **2.1% (623,699)** | **0.2% (1,155)** | **12.3% (3,697,430)** | **2.6% (12,630)** | **13.3% (3,998,585)** | **37.3% (183,672)** | **3.9% (1,179,553)** | **16.6% (81,627)** |
| Best PRS‡ | 70.4 | 1.6% (469,655) | 0.2% (887) | 10.8% (3,239,529) | 2.3% (11,323) | 12.9% (3,888,273) | 35.5% (174,598) | 3.6% (1,070,425) | 14.8% (72,613) |
| Classical risk factors + Best PRS‡ | 71.6 | 2.4% (720,155) | 0.3% (1,315) | 13.1% (3,948,851) | 2.7% (13,277) | 13.6% (4,077,750) | 38.5% (189,433) | 4.2% (1,250,309) | 17.8% (87,352) |
| Best PRS + Density‡,§ | 72.7 | 3.3% (991,883) | 0.4% (1,751) | 15.3% (4,582,391) | 3.0% (14,836) | 14.1% (4,238,237) | 41.2% (202,646) | 4.7% (1,412,975) | 20.6% (101,200) |
| **All risk factors‡,§** | **73.7** | **4.3% (1,297,046)** | **0.5% (2,225)** | **17.3% (5,208,326)** | **3.3% (16,314)** | **14.4% (4,339,033)** | **43.4% (213,683)** | **5.1% (1,545,326)** | **23.1% (113,621)** |

§ Odds ratio for mammographic breast density is adjusted for PRS

**Figure S1.**
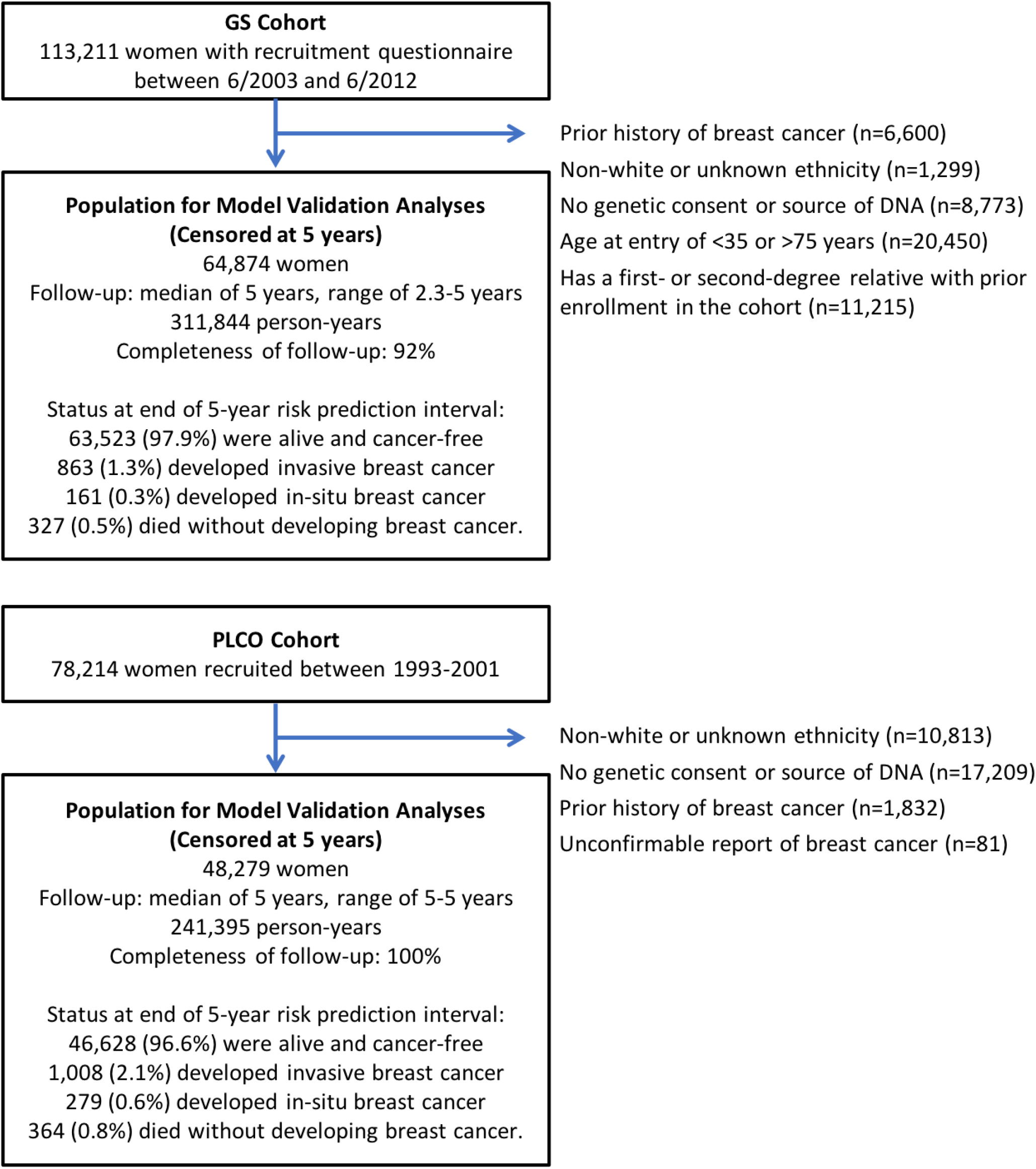
Study design of validation cohorts

**Figure S2.**
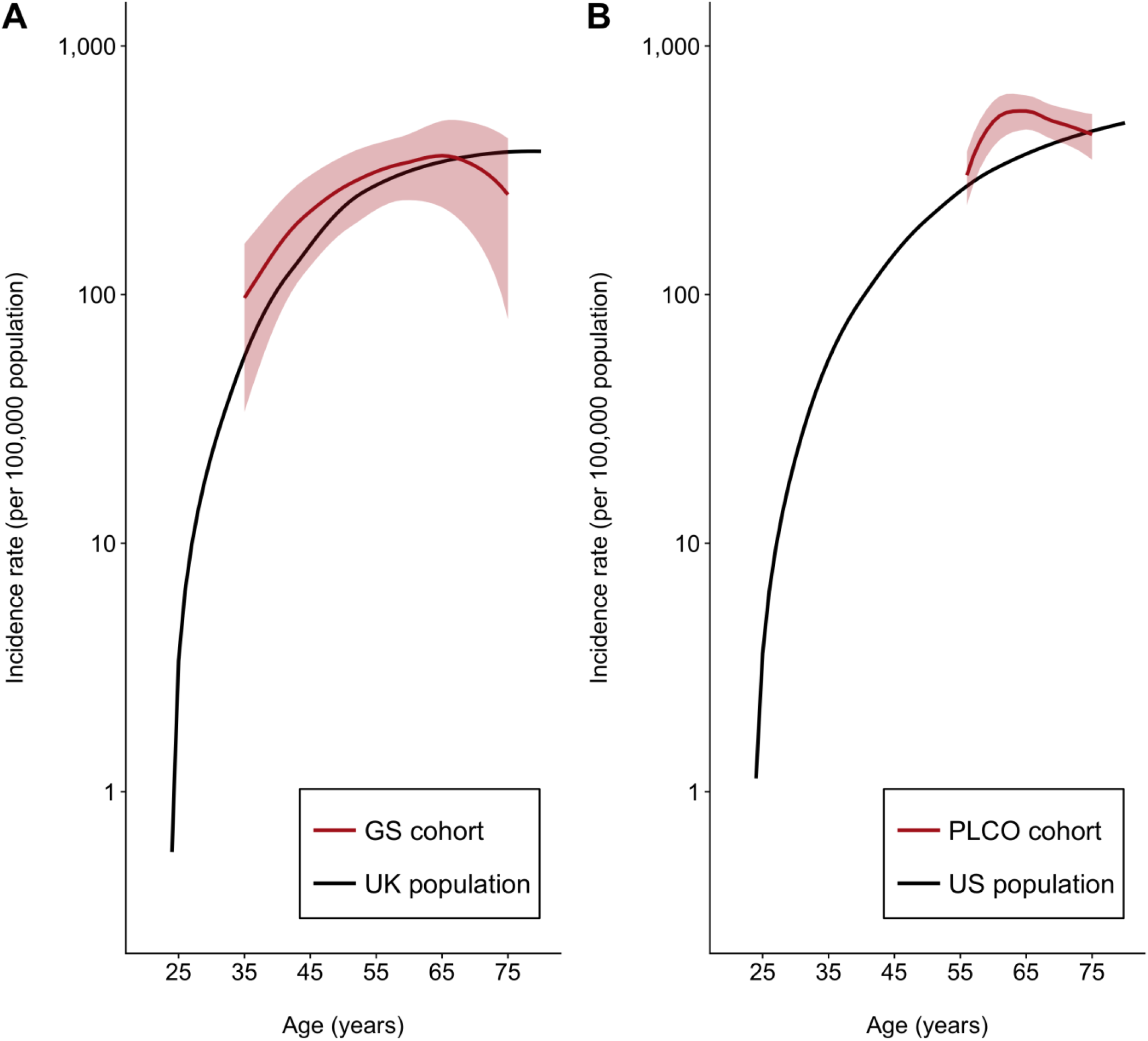
Breast cancer incidence rates in the validation cohort compared to the general population **(A)** Breast cancer incidence rates in the Generations Study (GS) cohort compared to the general UK population (ONS, 2006-2010). **(B)** Breast cancer incidence rates in the Prostate, Lung, Colorectal and Ovarian (PLCO) Cancer Screening Trial cohort compared to the general US population (SEER, 2010-2012).

**Figure S3A.**
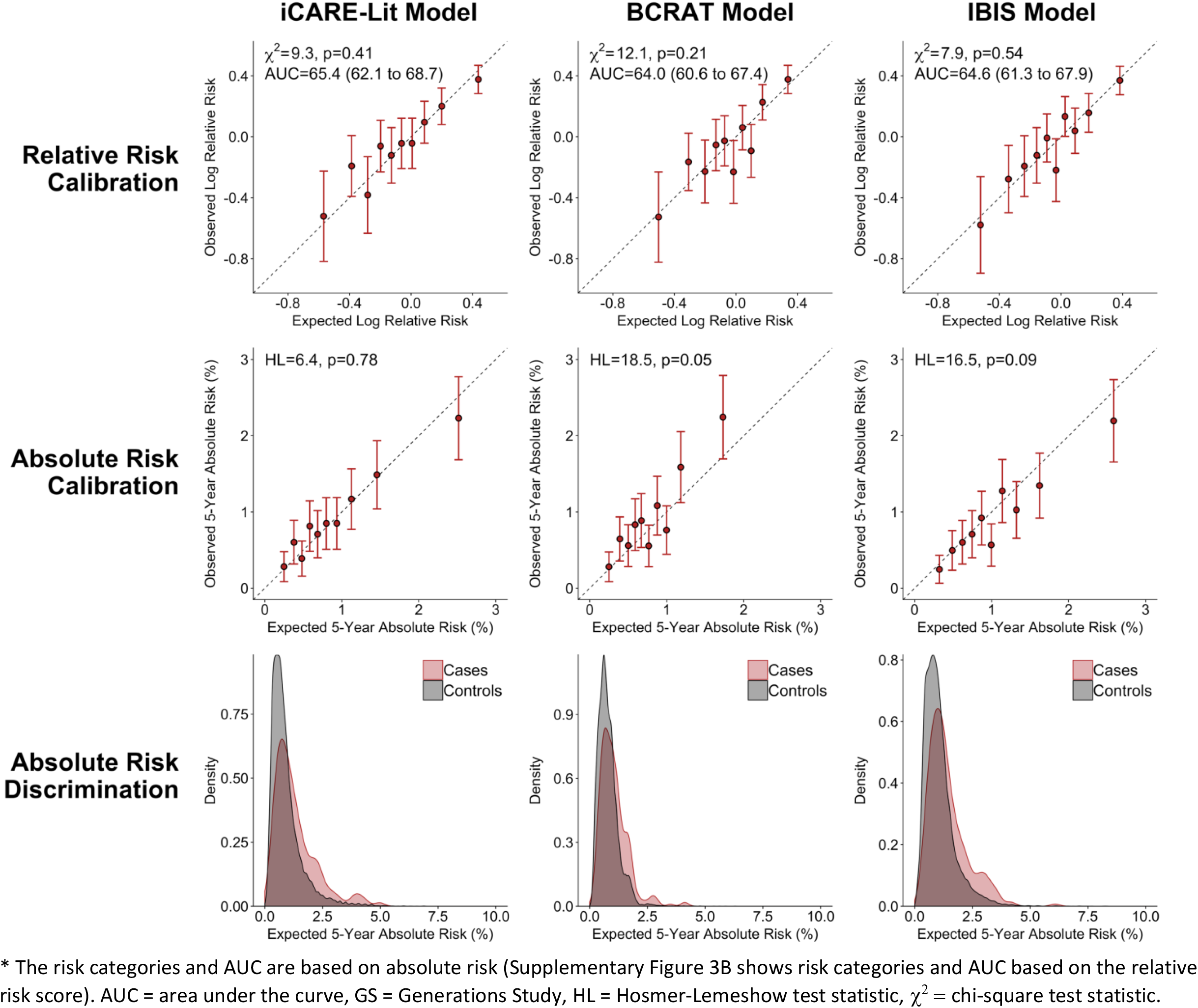
Calibration and discrimination, with risk categories based on **absolute risk**, of breast cancer risk prediction models in the **GS cohort among women less than 50 years of age**\*

**Figure S3B.**
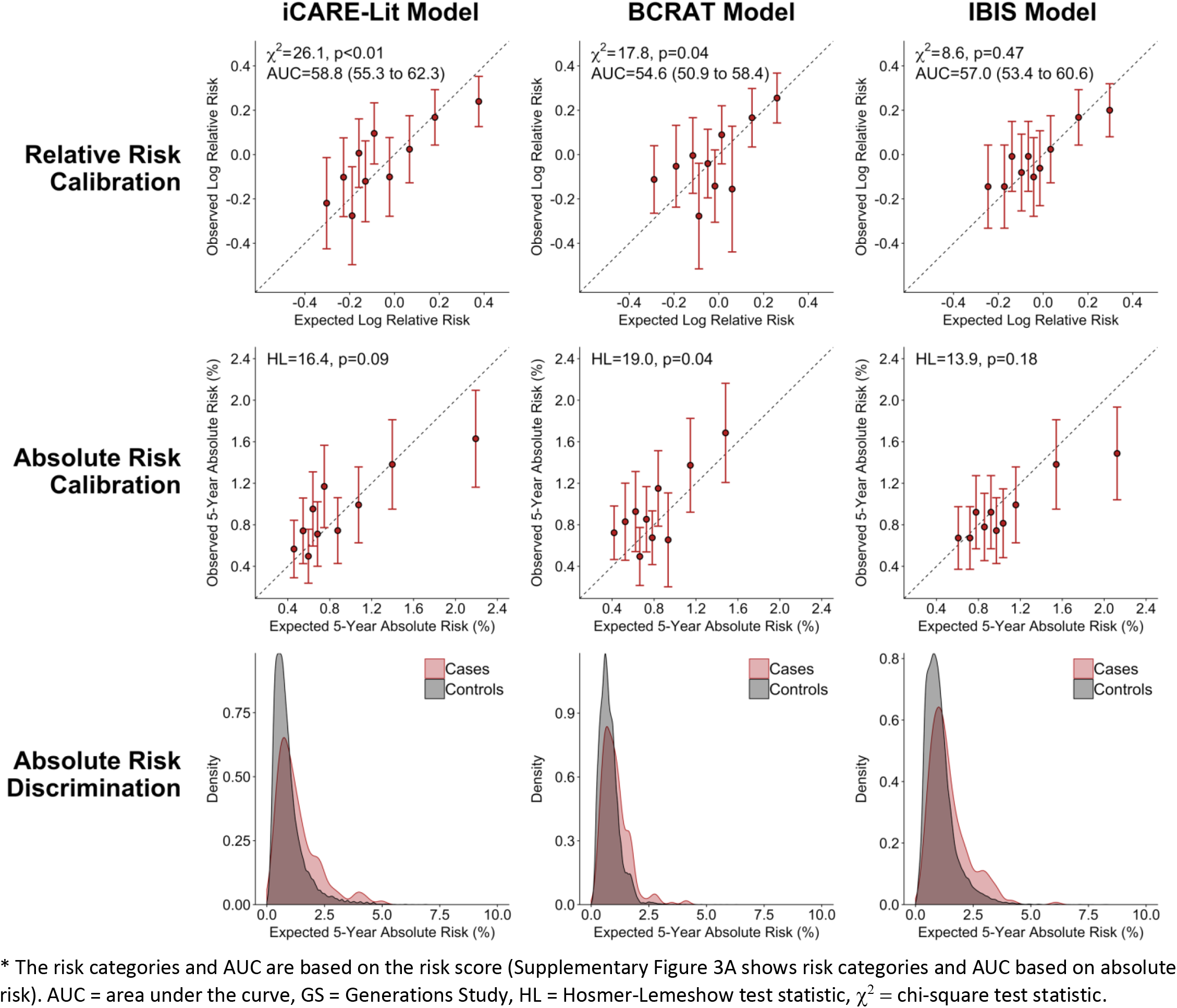
Calibration and discrimination, with risk categories based on the **relative risk score**, of breast cancer risk prediction models in the **GS cohort among women less than 50 years of age**\*

**Figure S4A.**
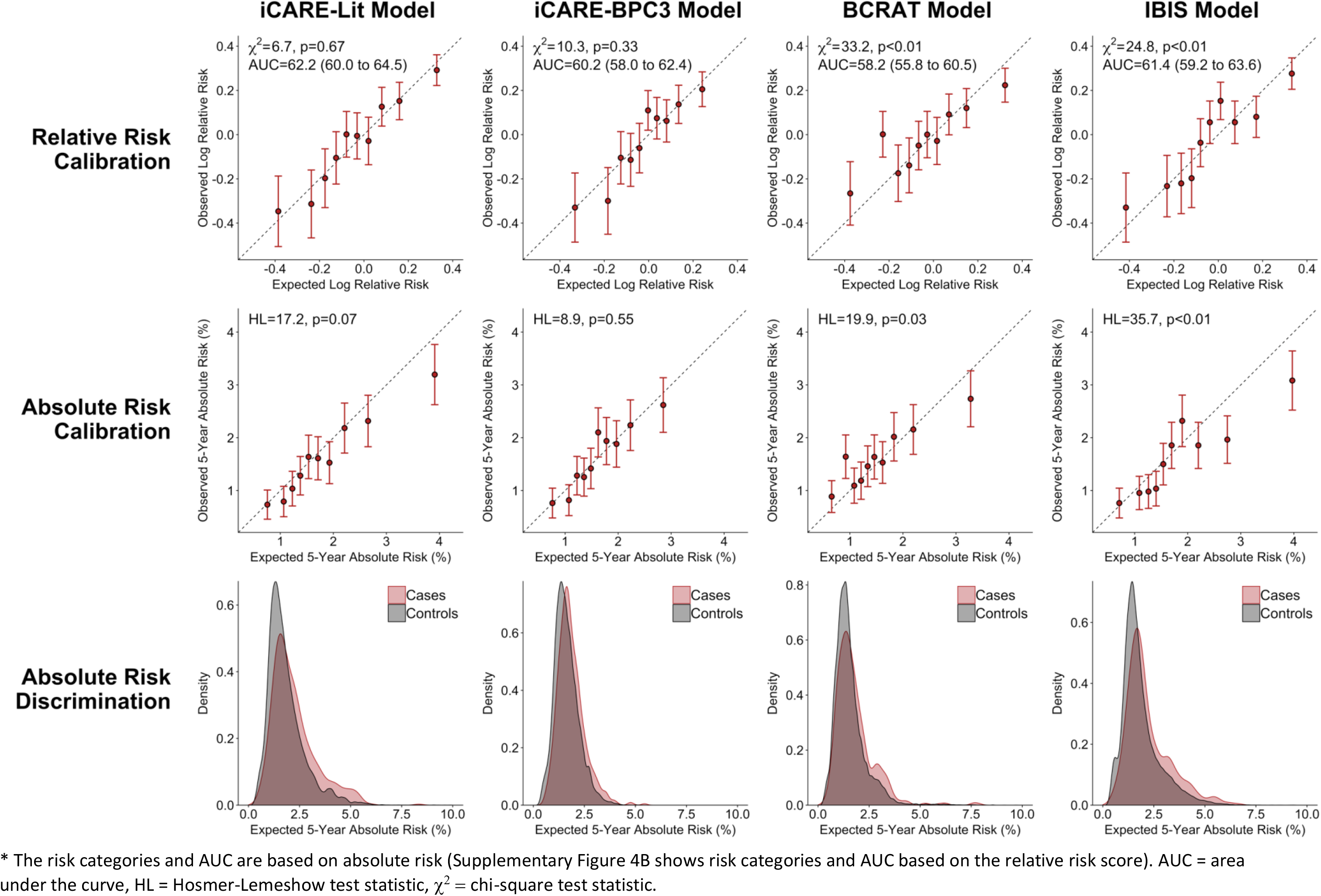
Calibration and discrimination, with risk categories based on **absolute risk**, of breast cancer risk prediction models in the **GS cohort among women 50 years of age or greater**\*

**Figure S4B.**
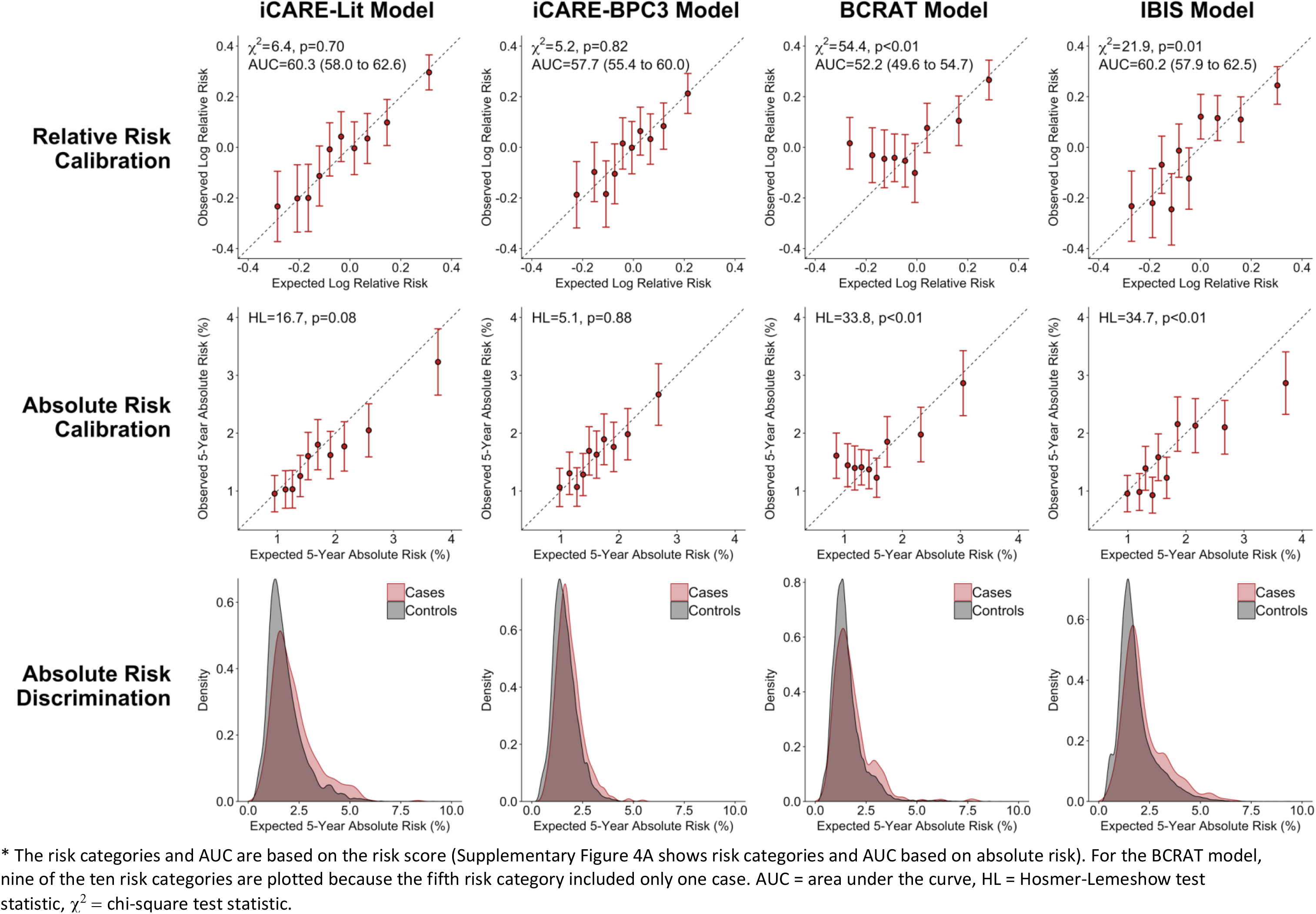
Calibration and discrimination, with risk categories based on the **relative risk score**, of breast cancer risk prediction models in the **GS cohort among women 50 years of age or greater**\*

**Figure S5A.**
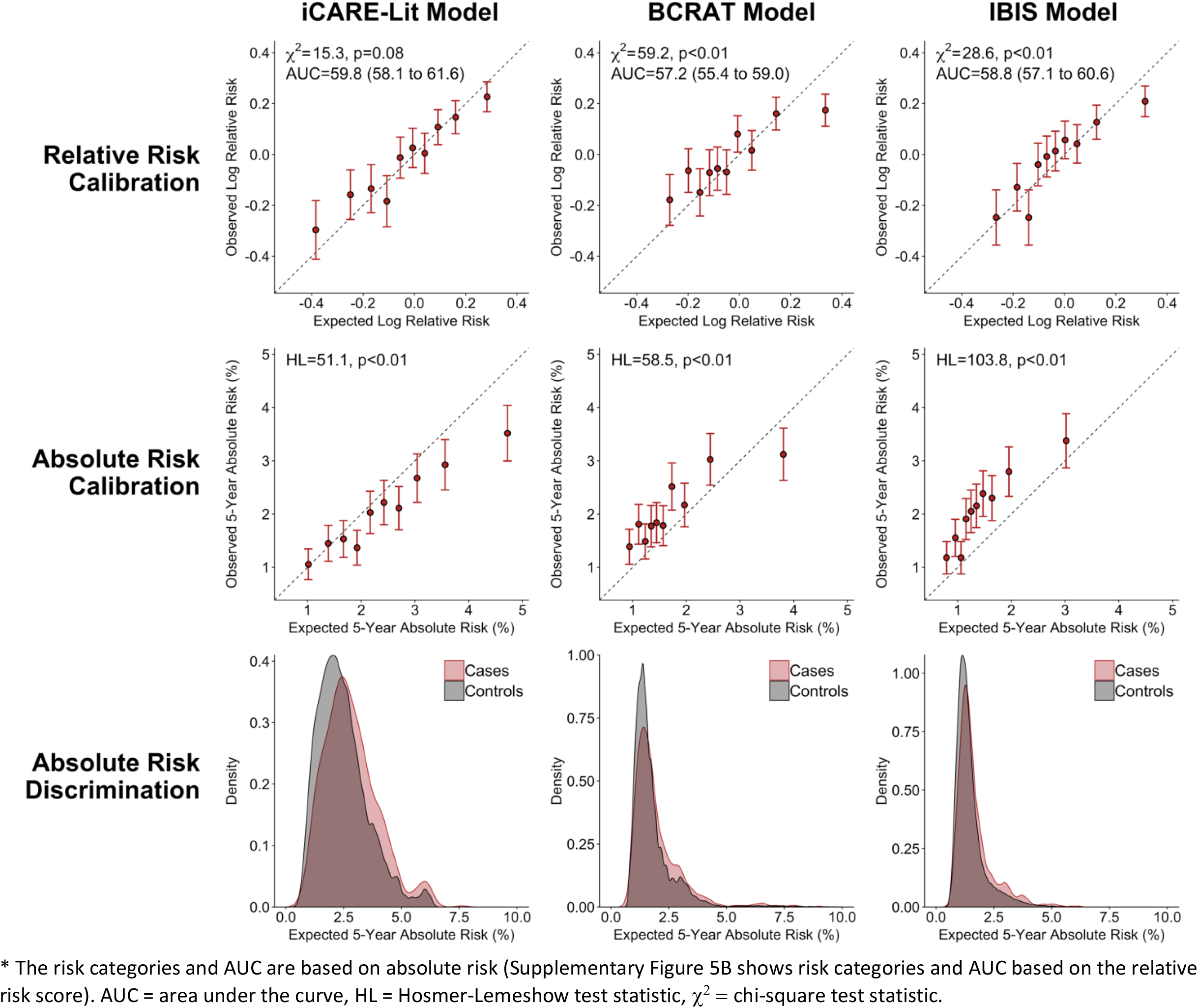
Calibration and discrimination, with risk categories based on **absolute risk**, of breast cancer risk prediction models in the **PLCO cohort among women 50 years of age or greater**\*

**Figure S5B.**
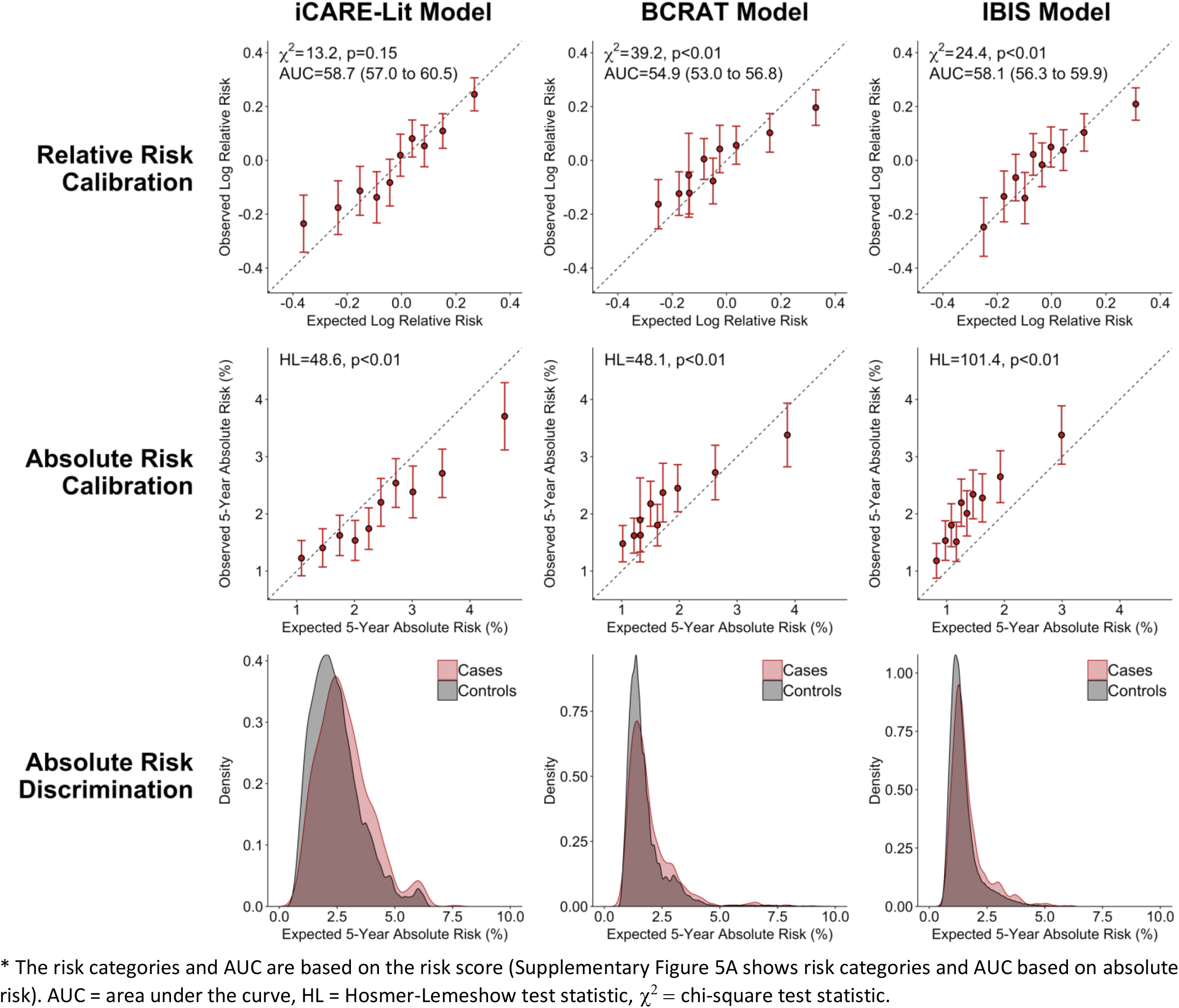
Calibration and discrimination, with risk categories based on the **relative risk score**, of breast cancer risk prediction models in the **PLCO cohort among women 50 years of age or greater**\*

## Notes

The authors have no conflicts of interest to disclose.

